# Sex Differences in the Brain’s White Matter Microstructure during Development assessed using Advanced Diffusion MRI Models

**DOI:** 10.1101/2024.02.02.578712

**Authors:** Sebastian M. Benavidez, Zvart Abaryan, Gaon S. Kim, Emily Laltoo, James T. McCracken, Paul M. Thompson, Katherine E. Lawrence

## Abstract

Typical sex differences in white matter (WM) microstructure during development are incompletely understood. Here we evaluated sex differences in WM microstructure during typical brain development using a sample of neurotypical individuals across a wide developmental age (N=239, aged 5-22 years). We used the conventional diffusion-weighted MRI (dMRI) model, diffusion tensor imaging (DTI), and two advanced dMRI models, the tensor distribution function (TDF) and neurite orientation dispersion density imaging (NODDI) to assess WM microstructure. WM microstructure exhibited significant, regionally consistent sex differences across the brain during typical development. Additionally, the TDF model was most sensitive in detecting sex differences. These findings highlight the importance of considering sex in neurodevelopmental research and underscore the value of the advanced TDF model.

## I. Introduction

Prior diffusion-weighted magnetic resonance imaging (dMRI) work has extensively characterized sex differences in the adult human brain using multiple dMRI models, providing evidence for subtle sex differences across the brain’s white matter (WM) for multiple dMRI metrics [1, 2]. However, less work has examined typical WM sex differences during development. Underscoring the importance of studying developmental sex differences, childhood through emerging adulthood is a period of substantial physiological and neural change – including in the brain’s WM microstructure [3, 4]. WM microstructure is implicated in multiple neuropsychiatric disorders that begin to manifest during adolescence and exhibit sex differences in their prevalence or presentation, including depression, bipolar disorder, and schizophrenia [5–7]. Understanding sex differences in WM microstructure during typical development may thus offer an important foundation for studying the neural mechanisms of such neuropsychiatric conditions.

Most studies of typical microstructural sex differences in individual WM tracts or regions during development have been constrained by the limitations of the diffusion model used, small sample sizes, and/or narrow age ranges. Most dMRI studies use the conventional diffusion tensor imaging (DTI) model [8]. However, the DTI model cannot accurately model complex fiber configurations, such as fiber crossing, due to its strict Gaussianity and use of a single tensor [9]. Most studies of regional WM sex differences in typical development examined fewer than 100 participants; existing larger studies have focused on narrow age ranges or early developmental periods that do not allow for the assessment of how sex differences may evolve between childhood and emerging adulthood [10– 12].

Diffusion models beyond the conventional DTI model can capture the underlying microenvironment more accurately and enable additional insights into WM microstructure. One such advanced diffusion model is the tensor distribution function (TDF), which uses a continuous mixture of Gaussian distributions to model diffusion and assigns weights to tensors based on their contribution to describing the diffusion in the voxel [13, 14]. TDF can also be computed from single-shell dMRI, making it well-suited for archival single-shell datasets and for developmental samples that require shorter scan acquisitions than adult samples. Prior work using TDF in Alzheimer’s disease, cognitive impairment, and normative aging has also shown that TDF can capture WM differences with greater sensitivity than DTI [2, 14, 15]. However, no prior work has used TDF to characterize WM microstructure in typical development. Advanced biophysical diffusion models are complementary to models such as TDF by reflecting additional properties of water diffusion that may correspond more closely with individual aspects of the cellular environment and thus allow for a more nuanced characterization of sex differences. One such biophysical model is the multi-compartment neurite orientation and dispersion density imaging (NODDI) model, which uses a Watson distribution to estimate the dispersion about the dominant orientation [16]. However, prior studies that have used NODDI to assess WM microstructure during typical development did not assess sex differences, focused on a selected region, or used very small samples [17–20]. TDF and NODDI have been widely used in multiple conditions, such as Alzheimer’s disease, traumatic brain injury, and aging, and have been shown to be more accurate in modeling WM microstructure compared to traditional DTI [2, 13-16, 21-22].

In this study, we used the dMRI models DTI, TDF, and NODDI to characterize WM sex differences in a sample of neurotypical individuals across a broad developmental age range. To ensure robust results, we conducted supplemental analyses that consider methodological and physiological factors important to developmental populations. We also assessed which dMRI model was the most sensitive to sex differences in developmental populations to inform future study design. As a whole, this comparatively large study (spanning this age range) is the first set of analyses to assess WM sex differences from childhood through emerging adulthood using the advanced dMRI models, TDF and NODDI.

## II. Methods

### A. Participants

The data used in this study is from the open-source Healthy Brain Network (HBN), an ongoing large-scale study across four data acquisition sites in the New York City area. The HBN initiative seeks to create a biobank of multimodal brain imaging and phenotypic data that captures a broad range of commonly encountered clinical psychopathology in childhood through emerging adulthood [23]. The HBN study was approved by the Chesapeake Institutional Review Board. Written informed consent was obtained from all participants ages 18 or older. For participants younger than 18, written informed consent was obtained from their legal guardian and written assent from each participant.

For the current analyses, we included neuroimaging and demographic data from neurotypical participants available through Release 10. As the purpose of the current study was to understand typical sex differences in white matter microstructure, included subjects were required to have completed a full clinical evaluation and received no diagnosis (i.e., neurotypical) and have a complete dMRI scan that passed quality control, including not having excessive head motion (see *MRI Acquisition and MRI Processing*). Sex was defined as the biological sex of the participant. Our final sample consisted of 239 neurotypical subjects between the ages of 5 to 22 years old (46.0% female; **Table 1**).

**Table 1:** Sample characteristics

|  | Boys | Girls | Boys vs. Girls |
| --- | --- | --- | --- |
| Sample Size | 129 | 110 | - |
| Age (years) | 11.44 ± 3.80 | 10.93 ± 4.21 | 0.34 |
| PPS Category | - | - | 0.01 |
| Pre-pubertal to early-pubertal | 52 | 29 | - |
| Mid-pubertal to post-pubertal | 27 | 37 | - |
| Head motion (mm) | 0.30 ± 0.22 | 0.29 ± 0.20 | 0.56 |

This table presents key demographic and clinical characteristics of the sample by sex. The age and head motion of participants are reported as mean ± standard deviation. ‘PPS’ denotes Peterson Puberty Scale staging, a measure used for assessing pubertal development. In the last column, p-values indicate the statistical significance of the differences observed between boys and girls.

Males and females in our sample did not significantly differ in age (*p*=0.16) or mean relative head motion (*p*=0.86). As expected, females were significantly more pubertally advanced than males (*p*=0.01) in the N=141 subset of participants with self-reported Peterson Puberty Scale (PPS) questionnaire scores [24, 25]. We thus included categorical puberty stage (pre-/early pubertal vs. mid-/late/post-pubertal) as a covariate in supplemental analyses to account for the unequal distribution of pubertal stages in our sample.

### B. MRI Acquisition

dMRI scans were acquired across four Siemens scanners in HBN: three Siemens 3T scanners (two Prismas and one Trio: 1.8 mm isotropic voxel size, 104 x 104 matrix, 72 slices, TR = 3320 ms, TE = 100.2 ms, multiband acceleration factor = 3), and one 1.5T scanner (Avanto: 2.0 mm isotropic voxel size, 96 x 96 matrix, 72 slices, TR = 4500 ms, TE = 93.8 ms, multiband acceleration factor = 3). All dMRI scans consisted of one *b* = 0 s/mm^2^ (*b*_*0*_) volume and 64 diffusion encoding directions for two diffusion-weighted shells: 64 directions at *b* = 1000 s/mm^2^ and 64 directions at *b* = 2000 s/mm^2^.

### C. MRI Processing

All raw T1-weighted and dMRI scans were visually inspected for quality assurance. Each subject’s dMRI scan was denoised using the Marcenko-Pastur principal component analysis (PCA) algorithm in DIPY to enhance the signal-to-noise ratio [26]. To correct for susceptibility artifacts in the dMRI images, we first generated an estimate of the susceptibility-induced off-resonance field calculated from a pair of reverse-phase encoded *b*_*0*_ images [27] using FSL’s *topup* [28]; for six participants who did not have a reverse-phase encoded *b*_*0*_ image, the off-resonance field was generated using the previously validated method, Synb0-DisCo, to create a synthetic *b*_*0*_ image derived from the dMRI and T1-weighted image [29]. FSL’s *eddy_cuda* then used this field to correct for susceptibility distortions, while also correcting for head motion and eddy currents, including rotating the *b*-matrix. We then corrected for bias field artifacts in the dMRI scans using ANTs’ *N4BiasFieldCorrection* algorithm [30], as implemented in MRtrix3’s *dwibiascorrect* [31, 32].

We employed three distinct diffusion reconstruction models to derive eight relevant metrics. Using FSL’s *dtifit*, we fit the traditional DTI model, which generates a single tensor, to derive measures of fractional anisotropy (FA^DTI^: indicative of non-uniform diffusion), mean diffusivity (MD: overall magnitude of diffusion), axial diffusivity (AD: diffusion in the principal direction), and radial diffusivity (RD: diffusion in the orthogonal directions) [8]. Using publicly available code (https://git.ini.usc.edu/ibagari/TDF), we fit the probabilistic TDF model, which models diffusion with a continuous mixture of Gaussians, to generate a measure of FA^TDF^ that is similar to FA^DTI^ but more accurately describes diffusion in areas of crossing fibers [13, 14]. For DTI and TDF, only the lower *b*-value shell, *b* = 1000 s/mm^2^, was used in the reconstruction, to be consistent with previous literature [2, 12, 33]. We fit the multi-compartment NODDI model using the Dmipy Toolbox [34], which models the dispersion of a single bundle using a Watson distribution [16]. Using this model, we derived measures of orientation dispersion index (ODI: orientation dispersion of the bundle), intracellular volume fraction (ICVF: restricted diffusion), and isotropic volume fraction (ISOVF: isotropic Gaussian diffusion). All diffusion-weighted shells were used to reconstruct the NODDI model.

To generate white matter summary measures for each region of interest (ROI), we used the ENIGMA-DTI protocol (http://enigma.ini.usc.edu/protocols/dti-protocols) [35]. Briefly, FSL’s tract-based spatial statistics (TBSS) [36] was used together with the ENIGMA-DTI template to skeletonize the FA^DTI^ image; the projection used for FA^DTI^ was then used to skeletonize all other dMRI metrics. The average of each diffusion metric in the skeleton was then calculated for 25 bilateral deep WM ROIs from the Johns Hopkins University (JHU) DTI atlas [37]: the corpus callosum (CC), genu of CC (GCC), body of CC (BCC), splenium of CC (SCC), fornix (FX), corticospinal tract (CST), internal capsule (IC), anterior limb of IC (ALIC), posterior limb of IC (PLIC), retrolenticular part of IC (RLIC), *corona radiata* (CR), anterior CR (ACR), superior CR (SCR), posterior CR (PCR), posterior thalamic radiation (PTR), *sagittal stratum* (SS), external capsule (EC), cingulum (cingulate gyrus) (CGC), cingulum (hippocampus) (CGH), fornix/*stria terminalis* (FX/ST), superior longitudinal fasciculus (SLF), superior fronto-occipital fasciculus (SFO), *tapetum* (TAP), uncinate fasciculus (UNC), and whole white matter average (full WM). To adjust for inter-scanner variability, we used ComBat (neuroCombat; [https://github.com/Jfortin1/neuroCombat]) to harmonize subjects’ ROI data within each dMRI metric while preserving variance due to age and sex [38, 39]. All image pre-processing and processing steps were checked visually for quality assurance, and subjects with excessive head motion were excluded.

### D. Statistics

Our primary analyses used a fixed-effects linear regression to assess sex differences in WM microstructure (main effect of sex), with additional analyses testing the interaction between sex and age (sex × age and sex × age^2^) to determine if age modified the effect of sex; all analyses also included demeaned age and age^2^ terms as nuisance covariates. We report effect sizes as the standardized beta, as we included nuisance covariates in our regression and standardized betas also allow for comparability with previous WM sex differences studies [10, 40]. A false discovery rate (FDR) [41] of 5% was used to correct for multiple comparisons across ROIs. Significance of all results were determined as *q* < 0.05 after FDR correction. Supplemental analyses were completed to determine if our results remained significant when including mean relative head motion or pubertal stage as additional nuisance covariates, and when using the alternative scanner harmonization method ComBat-GAM (NeuroHarmonize’s harmonizationLearn) [42]. ComBat-GAM removes batch effects between sites by fitting a generalized additive model (GAM) with a penalized nonlinear term to describe age effects.

## III. Results

Our analyses investigating the main effect of sex revealed significant microstructure differences between boys and girls in multiple WM ROIs across the brain (**Fig. 1–5**). As a whole, the DTI model showed that boys had reduced fractional anisotropy and greater diffusivity when compared to girls. Boys displayed significantly lower FA^DTI^ in the SS than girls, on average. Boys also exhibited significantly greater MD than girls in the full WM and in multiple ROIs; the largest effect size was observed in the SS, with similar effect sizes across the EC, UNC, PTR, SLF, CR, ACR, and PCR. For AD, boys displayed significantly higher values compared to girls in the SLF, SS, and PCR, with similar effect sizes across these ROIs. Compared to girls, boys also showed significantly greater RD in the SS. When using the TDF model, boys displayed significantly lower FA^TDF^ than girls, on average. The largest effect sizes for FA^TDF^ were in the SS, PCR, PTR, and FXST, with slightly smaller effect sizes in the CR, SCR, RLIC, TAP, SLF, and EC. For the NODDI measures, boys displayed significantly lower ICVF in the SS, TAP, and PTR compared to girls, with similar effect sizes across the ROIs. Boys also displayed significantly greater ISOVF than girls, on average, in the full WM and in multiple ROIs across the brain. The largest effect sizes for ISOVF were in the SS, EC, and CGH, with smaller effect sizes in the CR, ACR. For the ODI metric, no ROIs exhibited a significant main effect of sex. Altogether across the three dMRI models, multiple ROIs displayed significant sex differences: girls displayed higher anisotropy and restricted diffusion compared to boys, together with lower diffusivity and overall free diffusion than boys.

**Fig. 1.**
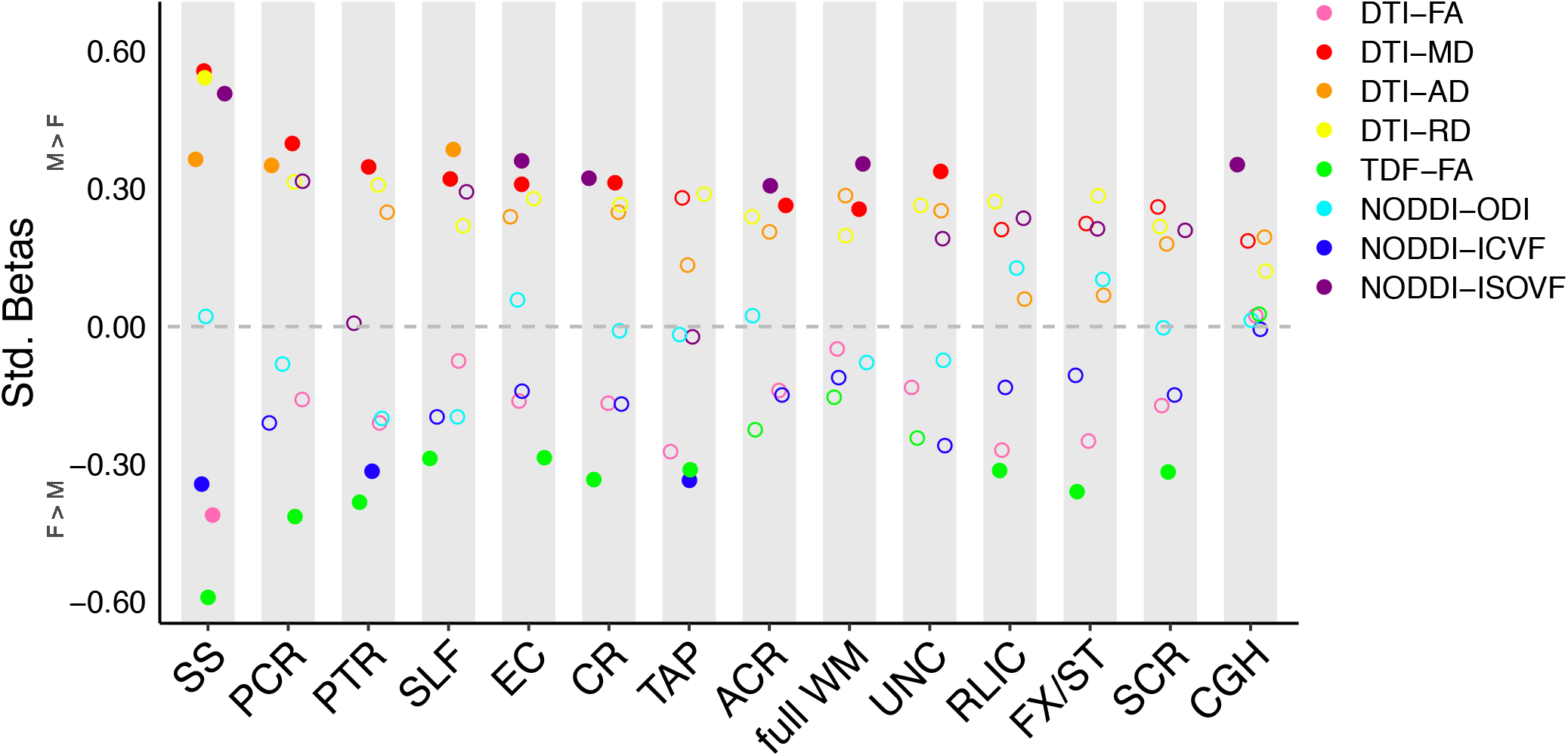
Effect sizes for the main effect of sex. ROIs are displayed if they exhibited a significant main effect of sex for at least one metric, with solid circles representing significant effects, and hollow circles representing non-significant effects. Positive effect sizes represent metrics that were greater in boys compared to girls, and negative effect sizes represent metrics that were greater in girls compared to boys.

**Fig. 2.**
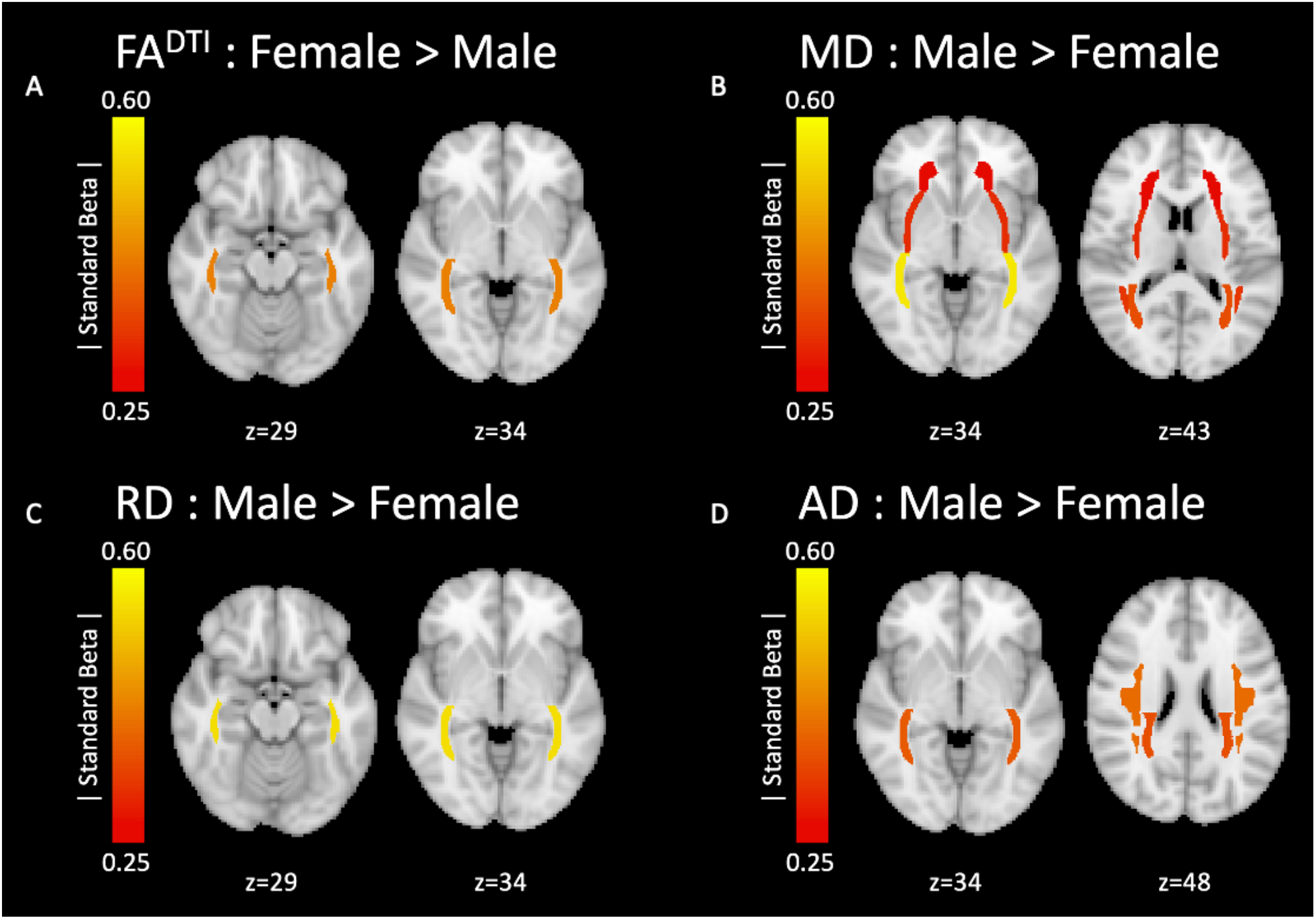
Magnitude of effect sizes for the main effect of sex on the DTI metrics. Significant ROIs are mapped onto the brain and colored according to the magnitude of the standard beta term for the main effect of sex. The yellow represents greater effect sizes and red represents smaller effect sizes. The title above each set of brain images indicates the direction of the effect. (A) the significant effect on FA^DTI^, (B) significant effects on MD, (C) significant effect on RD, and (D) significant effects on AD.

**Fig. 3.**
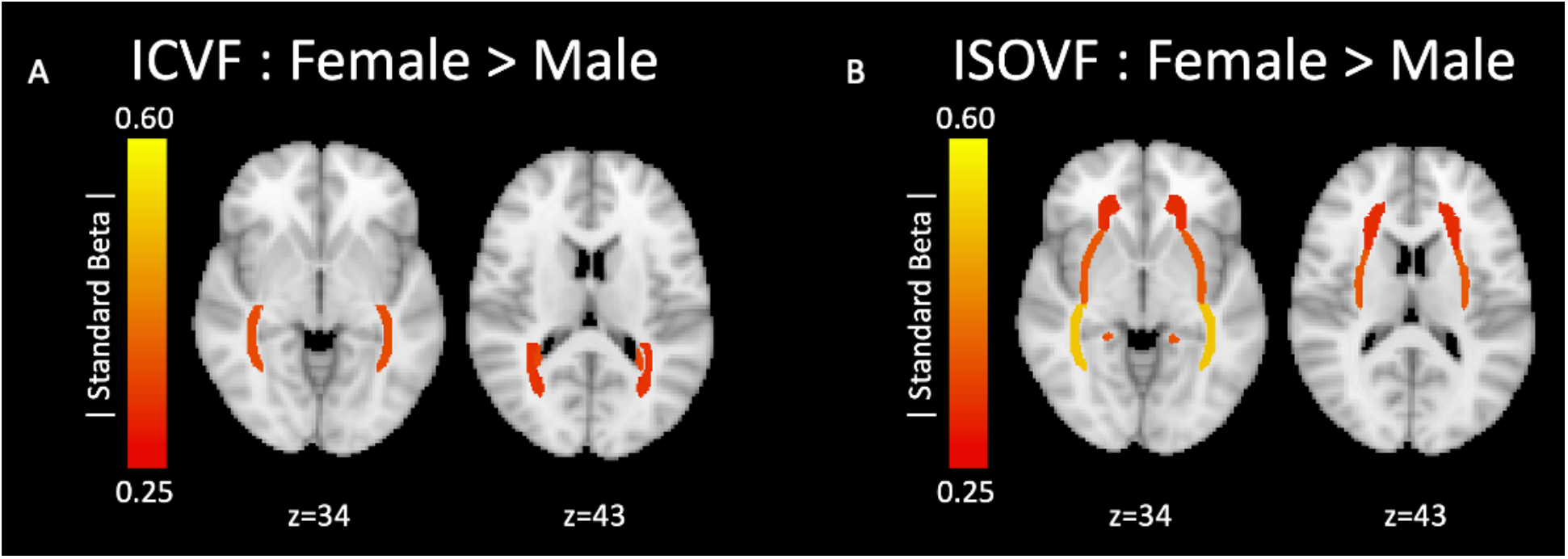
Magnitude of effect sizes for the main effect of sex on the NODDI metrics. Significant ROIs are mapped onto the brain and colored according to the magnitude of the standard beta term for the main effect of sex. The yellow represents greater effect sizes and red represents smaller effect sizes. The title above each set of brain images indicates the direction of the effect. (A) the significant effects on ICVF and (B) significant effects on ISOVF.

**Fig. 4.**
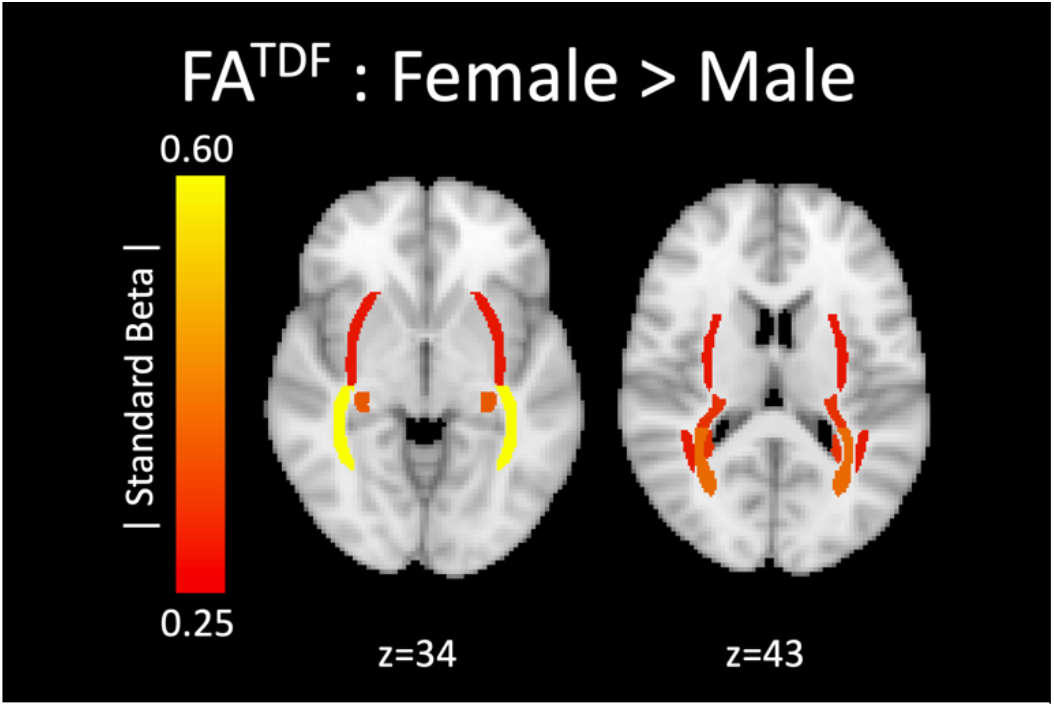
Magnitude of effect sizes for the main effect of sex on the FA^TDF^. Significant ROIs are mapped onto the brain and colored according to the magnitude of the standard beta term for the main effect of sex. The yellow represents greater effect sizes and red represents smaller effect sizes. The title above each set of brain images indicates the direction of the effect.

**Fig. 5.**
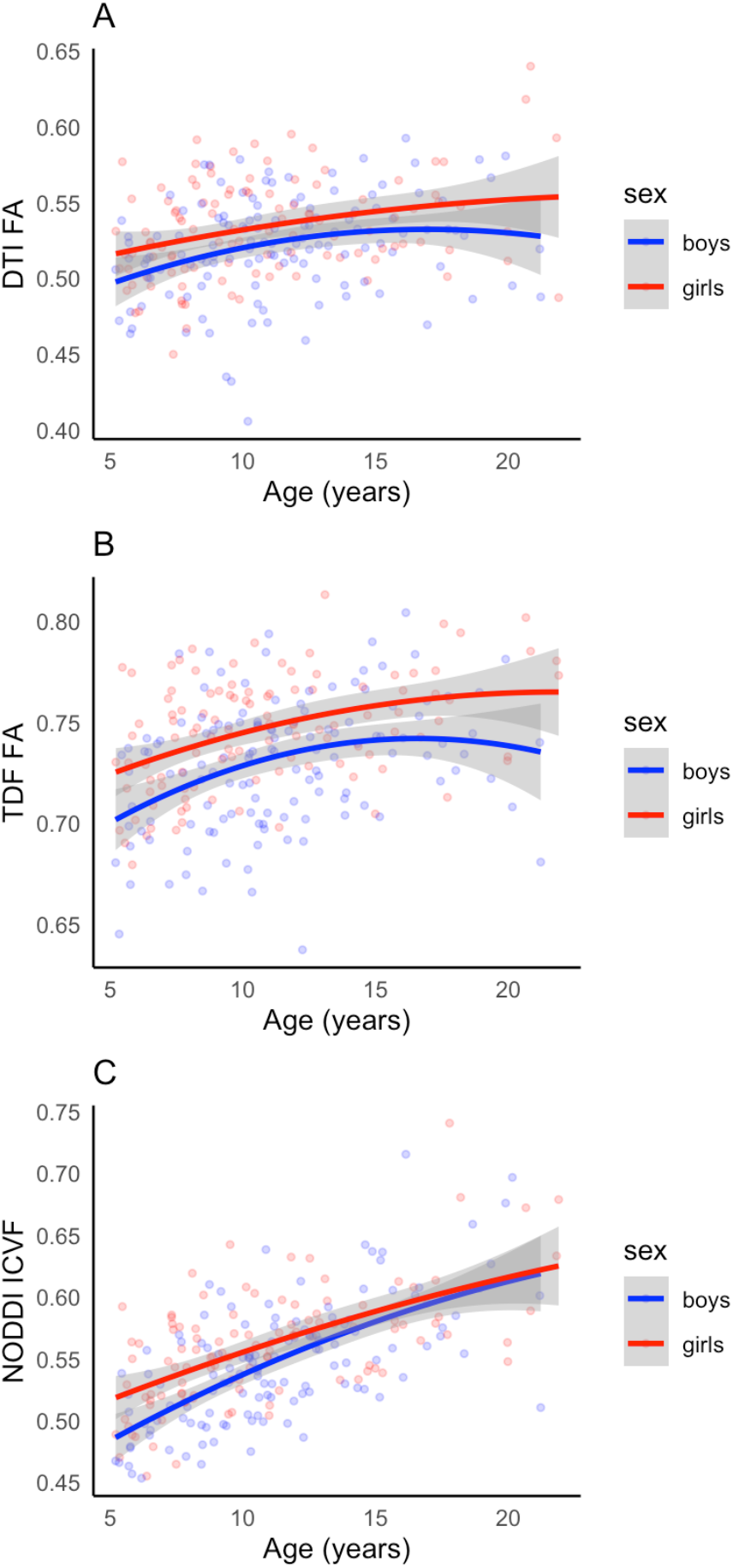
Plots displaying the distribution of boys and girls for dMRI metrics values of (A) FA^DTI^, (B) FA^TDF^, and (C) ICVF in the *sagittal stratum* plotted against age.

To consider the relative sensitivity of the different dMRI models to the main effect of sex (**Fig. 1**), we examined the number of significant ROIs in each of the three models. DTI captured significant sex effects in 9 ROIs, with the MD metric exhibiting the greatest sensitivity by capturing sex differences in all 9 significant ROIs. The TDF model captured significant effects on FA^TDF^ in 10 ROIs. Across all NODDI metrics, significant sex effects were observed in 8 ROIs; the most sensitive metric, ISOVF, captured significant effects in 6 ROIs. In sum, the TDF model detected significant differences between boys and girls in the greatest number of ROIs, although the number of significant ROIs was similar across all three dMRI models.

In supplemental analyses, we assessed the robustness of our significant sex difference results to multiple considerations, including scanner harmonization method, head motion, and pubertal development. As a whole, most of our sex difference findings remained significant. When using the harmonization method ComBat-GAM, the observed sex effects remained significant for all metrics in all ROIs. All sex difference results similarly remained significant when statistically covarying for mean relative head motion. When only considering participants with complete pubertal data and including pubertal stage as a nuisance covariate to account for the unequal distribution of pubertal stage in our sample, many of the metric and ROI combinations continued to exhibit significant sex differences. For the DTI measures, FA^DTI^ and RD in the SS remained significant for the main effect of sex. For AD, only the SLF remained significant. For MD, the SS, UNC, and PCR remained significant. TDF’s FA^TDF^ measure remained significant for the SS, PCR, CR, and EC. In the NODDI model, the SS remained significant for ICVF. For ISOVF, the SS, CGH, and full WM remained significant. In sum, multiple sex difference results remained significant in the supplemental analyses, with the most robust results seen in the SS, CR, and PCR.

Our analyses examining the interaction between sex and age demonstrated a significant interaction for the NODDI metric ODI in the SCR only. Visual inspection of the data revealed that boys display declining ODI between early childhood and emerging adulthood, whereas girls display increasing ODI from adolescence through emerging adulthood, such that girls exhibit greater ODI than boys by emerging adulthood. This interaction between sex and age remained significant in the supplemental analyses using ComBat-GAM and covarying for mean relative head motion, although it no longer quite attained significance when only including participants with complete pubertal data and covarying pubertal stage (*p*=0.055). In short, ODI exhibits a significant interaction between sex and age that is relatively robust.

## IV. Discussion

Here we used multiple advanced dMRI models to rigorously characterize sex differences in WM microstructure during typical development, including how such differences may vary between childhood and emerging adulthood. Neurotypical boys and girls exhibited significant microstructure differences in a range of deep WM regions across the brain when using the conventional model, DTI, as well as the advanced models, TDF and NODDI. The directionality of such sex differences was consistent across all WM regions assessed here. Compared to boys, girls exhibited greater FA^DTI^, FA^TDF^, and ICVF than boys, on average. Boys displayed higher MD, AD, RD, and ISOVF than girls, on average. These sex differences were observed in WM regions that included a mixture of projection, association, commissural, and limbic tracts. When considering how sex differences depended on age in our cross-sectional sample, the greatest sex differences in ODI were observed in emerging adulthood.

We also considered the relative sensitivity of the DTI, TDF, and NODDI models to sex effects on WM microstructure in our developmental sample. When examining sex differences across development, TDF detected significant sex differences in the greatest number of WM regions, followed by DTI and NODDI; however, the overall sensitivity of these three models to the main effect of sex was similar. When investigating how sex differences are modulated by age, NODDI was the only reconstruction model to capture a significant interaction between sex and age. Taken together, these findings suggest the utility of including advanced dMRI models when evaluating WM sex differences in developmental populations.

The current study has several strengths. The use of multiple dMRI models allows for a more thorough characterization of white matter sex differences, including the relative sensitivity of each model to these differences. Our use of the advanced model, TDF, also allows for greater compatibility with large-scale single-shell archival dMRI data, as well as newly collected single-shell dMRI scans in populations less likely to tolerate long multi-shell scans (e.g., infants) [43, 44]. Our results from the advanced multi-shell model NODDI may provide greater biological specificity than DTI or TDF by directly modeling multiple aspects of the cellular environment [16], although some recent studies suggest that the assumptions underlying the specificity of NODDI’s metrics may not always be met [45, 46]. Future studies should assess the generalizability of our findings by including additional dMRI models (e.g., restriction spectrum imaging) and using complementary dMRI analytic techniques beyond the TBSS method used here (e.g., tractography).

## V. Conclusion

We found widespread and regionally consistent sex effects on WM microstructure during development. These results expand on prior neurodevelopmental studies that used the DTI model to examine regional WM sex differences in smaller samples or narrower age ranges [10–12]. Furthermore, these findings provide the first robust characterization of regional sex differences in WM microstructure between childhood and emerging adulthood when using the advanced dMRI models, TDF and NODDI. In sum, our study provides an important reference for the future analysis of sex differences in typical development and adolescent-onset neuropsychiatric conditions.

